# Investigations of dimethylglycine (DMG), glycine betaine and ectoine uptake by a BCCT family transporter with broad substrate specificity in *Vibrio* species

**DOI:** 10.1101/2020.05.29.123752

**Authors:** Gwendolyn J. Gregory, Anirudha Dutta, Vijay Parashar, E. Fidelma Boyd

## Abstract

Fluctuations in osmolarity are one of the most prevalent stresses to which bacteria must adapt, both hypo- and hyper-osmotic conditions. Most bacteria cope with high osmolarity by accumulating compatible solutes (osmolytes) in the cytoplasm to maintain the turgor pressure of the cell. *Vibrio parahaemolyticus*, a halophile, utilizes at least six compatible solute transporters for the uptake of osmolytes: two ABC family ProU transporters and four betaine-carnitine-choline transporter (BCCT) family transporters. The full range of compatible solutes transported by this species has yet to be determined. Using an osmolyte phenotypic microarray plate for growth analyses, we expanded known osmolytes used by *V. parahaemolyticus* to include N-N dimethylglycine (DMG) amongst others. We showed that *V. parahaemolyticus* requires a BCCT transporter for DMG uptake, carriers that were not known to transport DMG. Growth pattern analysis of four triple-*bccT* mutants, possessing only one functional BCCT, indicated that BccT1 (VP1456), BccT2 (VP1723), and BccT3 (VP1905) transported DMG, which was confirmed by functional complementation in *E. coli* strain MKH13. BccT1 was unusual in that it could uptake both compounds with methylated head groups (glycine betaine (GB), choline and DMG) and cyclic compounds (ectoine and proline). Bioinformatics analysis identified the four coordinating residues for glycine betaine in BccT1. *In silico* modelling analysis demonstrated that glycine betaine, DMG, and ectoine docked in the same binding pocket in BccT1. Using site-directed mutagenesis, we showed that a strain with all four resides mutated resulted in loss of uptake of glycine betaine, DMG and ectoine. We showed three of the four residues were essential for ectoine uptake whereas only one of the residues was essential for glycine betaine uptake. Overall, we have demonstrated that DMG is a highly effective compatible solute for *Vibrio* species and have elucidated the amino acid residues in BccT1 that are important for coordination of glycine betaine, DMG and ectoine transport.

**Importance:** *Vibrio parahaemolyticus* possesses at least six osmolyte transporters, which allow the bacterium to adapt to high salinity conditions. In this study, we identified several novel osmolytes that are utilized by *V. parahaemolyticus*. We demonstrated that the compound dimethylglycine (DMG), which is abundant in the marine environment, is a highly effective osmolyte for *Vibrio* species. We determined that DMG is transported via BCCT-family carriers, which have not been shown previously to uptake this compound. BccT1 was a carrier for glycine betaine, DMG and ectoine and we identified the amino acid residues essential for coordination of these compounds. The data suggest that for BccT1, glycine betaine is more easily accommodated than ectoine in the transporter binding pocket.

## Introduction

In order to grow in high osmolarity environments, bacteria accumulate compounds called compatible solutes (osmolytes) within the cytoplasm of the cell, either via uptake or biosynthesis (1–4). These compounds balance the internal osmolarity with that of the environment and maintain the turgor pressure of the cell (3, 5, 6). Osmolytes also protect proteins, nucleic acids and other vital molecular machinery by increasing the hydration shell around these molecules (7). Osmolytes fall into several classes of compounds including: sugars (trehalose), polyols (glycerol, mannitol), free amino acids (proline, glutamine), amino acid derivatives (ectoine), and quarternary amines (glycine betaine (GB), carnitine) (4, 5, 8–12).

Biosynthesis of compatible solutes is energetically costly and therefore bacteria encode compatible solute transporters to scavenge available osmolytes from the environment (12–14). Compatible solute transporters include the multicomponent ATP Binding Cassette (ABC)-family such as ProU (*proVWX*) in *Escherichia coli* and OpuC (*proVWX*) in *Pseudomonas syringae* (15–17), and the single component betaine-carnitine-choline transporter (BCCT) family which are Na^+^ or H^+^ dependent. Members of the BCCT family include BetT, in *E. coli*, which transports choline with high affinity, and glycine betaine transporters in *Bacillus subtilis* (OpuD) and *Corynebacterium glutamicum* (BetP), among many others (18–21).

BCCTs are energized by sodium- or proton-motive force symport and are organized into 12 transmembrane (TM) segments (18, 20, 22, 23). Aromatic residues found in TM4 and TM8 make up the glycine betaine binding pocket in BCCT-family transporters examined to date. These residues are highly conserved in BCCTs that transport trimethylammonium compounds such as glycine betaine, L-carnitine and γ-butyrobetaine (19, 22, 24, 25). An additional tryptophan residue is present in TM8 just outside the binding pocket, and is thought to participate in coordination of substrates during conformational changes that occur during transport (24).

*Vibrio parahaemolyticus* is a halophilic bacterium that is found in marine and estuarine environments in association with plankton, fish and shellfish (26–31). *V. parahaemolyticus* is the leading bacterial cause of seafood-related gastroenteritis worldwide, frequently associated with consuming raw or undercooked seafood (28, 32). *Vibrio parahaemolyticus* can grow in a range of salinities and possesses four BCCTs encoded by *bccT1* (VP1456), *bccT2* (VP1723), *bccT3* (VP1905), and *bccT4* (VPA0356), and two ProUs encoded by *proVWX* (VP1726-VP1728) and *proXWV* (VPA1109-VPA1111) for the uptake of compatible solutes (33). In addition to compatible solute transporters, *V. parahaemolyticus* also possesses biosynthesis systems for compatible solutes ectoine, *ectABC-asp_ect* (VP1719-VP1722) and glycine betaine, *betIBA-proXWV* (VPA1112-VPA1114) whose expression is induced in high salinity (33). It was demonstrated that the expression of these biosynthesis systems in *Vibrio* species was under tight regulation, controlled by quorum sensing regulators OpaR and AphA, as well as CosR a global regulator of the osmotic stress response (34–37). We reported previously that in *V. parahaemolyticus bccT1*, *bccT3*, *bccT4*, and both *proU* operons, are responsive to salinity, with the exception of *bccT2* (37, 38). Both *bccT1* and *bccT3* and both *proU* operons are repressed in low salinity by CosR (37, 38). Studies have shown that BccT1 in *V. parahaemolyticus* had the broadest substrate specificity, transporting glycine betaine, choline, proline, and ectoine, while BccT2 and BccT3 transport glycine betaine, choline and proline, and BccT4 transports only choline and proline (**Fig. 1**) (38). Interestingly, the number of osmolyte transporters present among *Vibrio* species varies, which suggested differences in osmolytes utilized and osmotolerance (33). It was shown that *Vibrio alginolyticus* contained four BCCTs and two ProUs, *Vibrio harveyi* and *Vibrio splendidus* possessed six BCCTs and two ProUs whereas *Vibrio cholerae* and *Vibrio vulnificus* possessed one BCCT, a BccT3 homolog (33).

**Figure 1.**
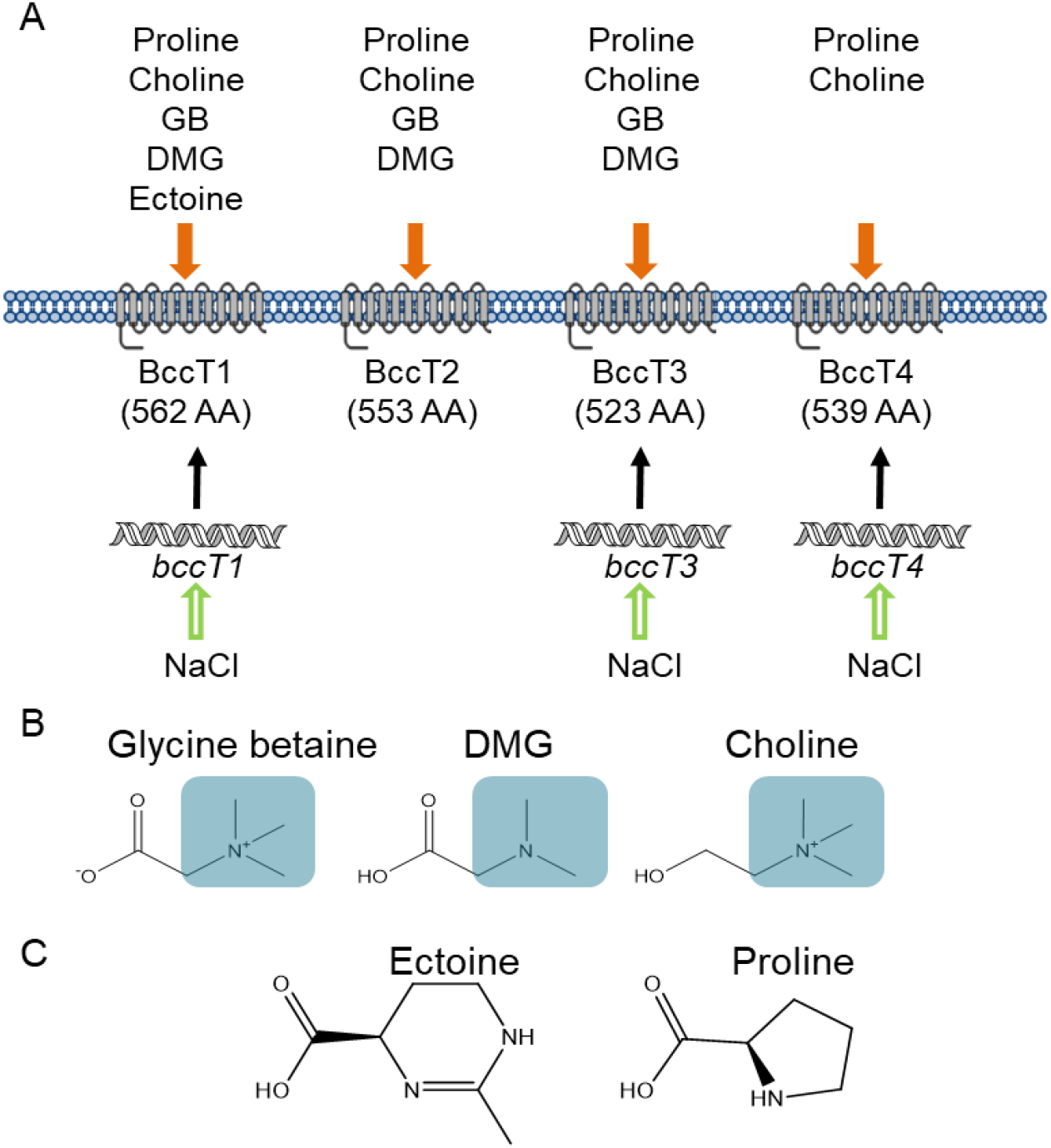
**(A)** BCCT transporters present in *V. parahaemolyticus* and their known substrates. *bccT1*, *bccT3*, and *bccT4* are induced by high salinity. Structures of BCCT substrates **(B)** with methylated headgroups highlighted in blue boxes or **(C)** cyclic compounds.

In this study, we examined the full range of osmolytes utilized by *V. parahaemolyticus* using an osmolyte phenotypic microarray plate, which identified over half a dozen potential osmolytes. We examined the ability of several different *Vibrio* species to utilize dimethylglycine, a compound previously not shown as an osmolyte in these species and one of the most effective osmolytes we identified. How *Vibrio* species uptake DMG is unknown. Therefore, we examined *V. parahaemolyticus* transport of DMG using several osmolyte transporter mutants. These analyses showed that the BCCT carriers were required for efficient DMG uptake, thus representing a new transporter family for the uptake of DMG. In *V. parahaemolyticus*, BccT1 has the broadest substrate uptake ability in terms of number and diversity of compounds (methylated head groups and cyclic compounds). Our *in silico* modelling analysis demonstrated that glycine betaine, DMG, and ectoine docked in the same binding pocket in BccT1. We investigated via mutagenesis and functional complementation the amino acid residues that are required for coordination of glycine betaine, DMG, and ectoine. This analysis describes for the first time the residues that coordinate DMG and ectoine in a BCCT family transporter.

## Results

### *V. parahaemolyticus* can utilize a wide range of compatible solutes

To determine the range of compatible solutes that can be utilized by *V. parahaemolyticus*, a Biolog 96-well PM9 osmolyte phenotypic microarray plate was used. Growth analyses was performed using a *V. parahaemolyticus* Δ*ectB* deletion mutant, which is unable to synthesize ectoine *de novo* and therefore has a growth defect in high salinity in the absence of exogenous osmolytes (33, 39). The Δ*ectB* mutant was grown in a Biolog 96-well PM9 osmolyte phenotypic microarray plate, which contains 23 unique osmolytes. A total of 14 of these osmolytes significantly rescue the growth of the Δ*ectB* mutant, which indicated that the substrate was transported and utilized by *V. parahaemolyticus* (**Fig. S1**). Previously unrecognized osmolytes for this species included N-N dimethylglycine (DMG), γ-amino-N-butyric acid (GABA), trimethylamine-N-oxide (TMAO), glutathione, dimethylsulfoniopropionate, MOPS, creatine, N-acetyl L-glutamine, and octopine, in addition to those already known to provide osmoprotection such as trehalose, β-glutamic acid, glycine betaine, ectoine and L-proline (**Fig. S1**).

To validate the phenotypic microarray analysis, we performed growth analyses with the Δ*ectB* mutant in the presence of compatible solutes DMG, GABA, or TMAO in in M9 minimal media supplemented with glucose and 6% NaCl (M9G 6%NaCl). The presence of exogenous DMG, GABA, or TMAO rescued the growth of the Δ*ectB* mutant, which confirmed *V. parahaemolyticus* can utilize these as compatible solutes (**Fig. 2**). Growth of the Δ*ectB* mutant was rescued by addition of exogenous DMG with a lag phase of less than one hour (**Fig. 2A**), which indicated that DMG is a very effective compatible solute for *V. parahaemolyticus*. The Δ*ectB* mutant was rescued to a greater extent with exogenous DMG as compared to GABA or TMAO (**Fig. 2B and 2C**).

**Figure 2.**
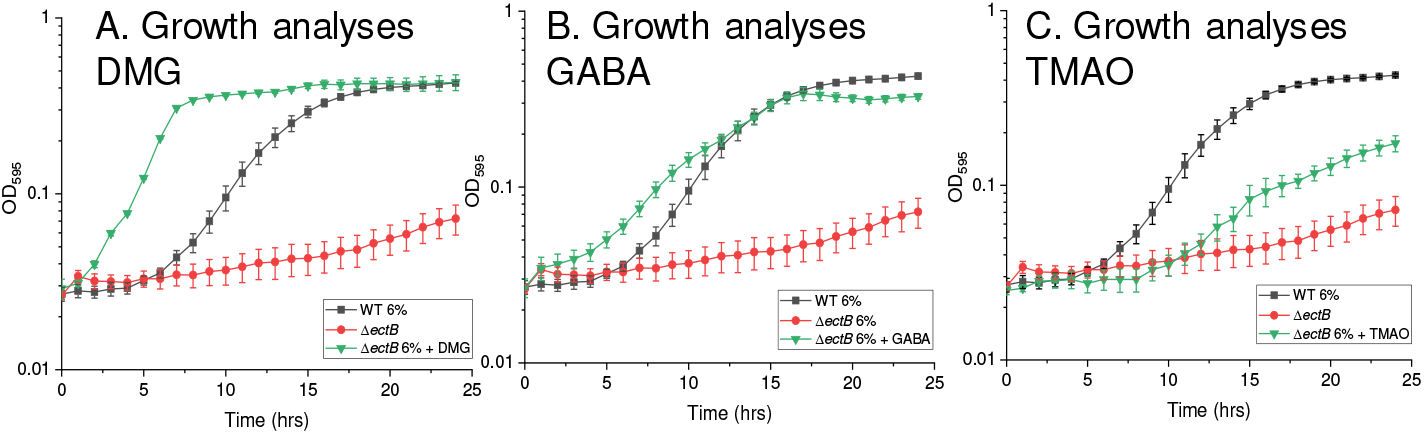
Growth analyses in M9G 6%NaCl with wild type and an *ectB* mutant strain. Media was supplemented with **(A)** DMG, **(B)** GABA, or **(C)** TMAO. Optical density (OD_595_) was measured every hour for 24 hours; the mean and standard error of at least two biological replicates are shown

### DMG is an important compatible solute for *Vibrio* species

To investigate whether DMG can also act as an osmolyte in other *Vibrio* species, we grew four species representing divergent clades of *Vibrio* in M9G 4% NaCl or 5%NaCl supplemented with and without DMG at 37C for 24 h. Growth of *V. harveyi, V. fluvialis*, *V. vulnificus* and *V. cholerae* were all rescued by exogenous DMG (**Fig. 3A-D**). In the absence of DMG, *V. harveyi* grew with a 9 h lag phase and a 3 h lag phase and a shorter doubling time in the presence of DMG (**Fig. 3A**), which indicated that DMG is a highly effective compatible solute for this species. Similarly, *V. fluvialis* grown in the presence of DMG had a reduced lag phase and a shorter doubling time (**Fig. 3B**), indicating DMG is an effective osmolyte. *V. vulnificus* is unable to grow in M9G 4%NaCl in the absence of DMG, but growth was rescued in the presence of DMG with a 10 h lag phase (**Fig. 3C**), indicating this is a highly effective osmolyte for this species. In the presence of DMG the growth rate of *V. cholerae* was significantly increased (**Fig. 3D**). These data indicated that all *Vibrio* species tested utilize DMG as an osmoprotectant, with varying degrees of effectiveness.

**Figure 3.**
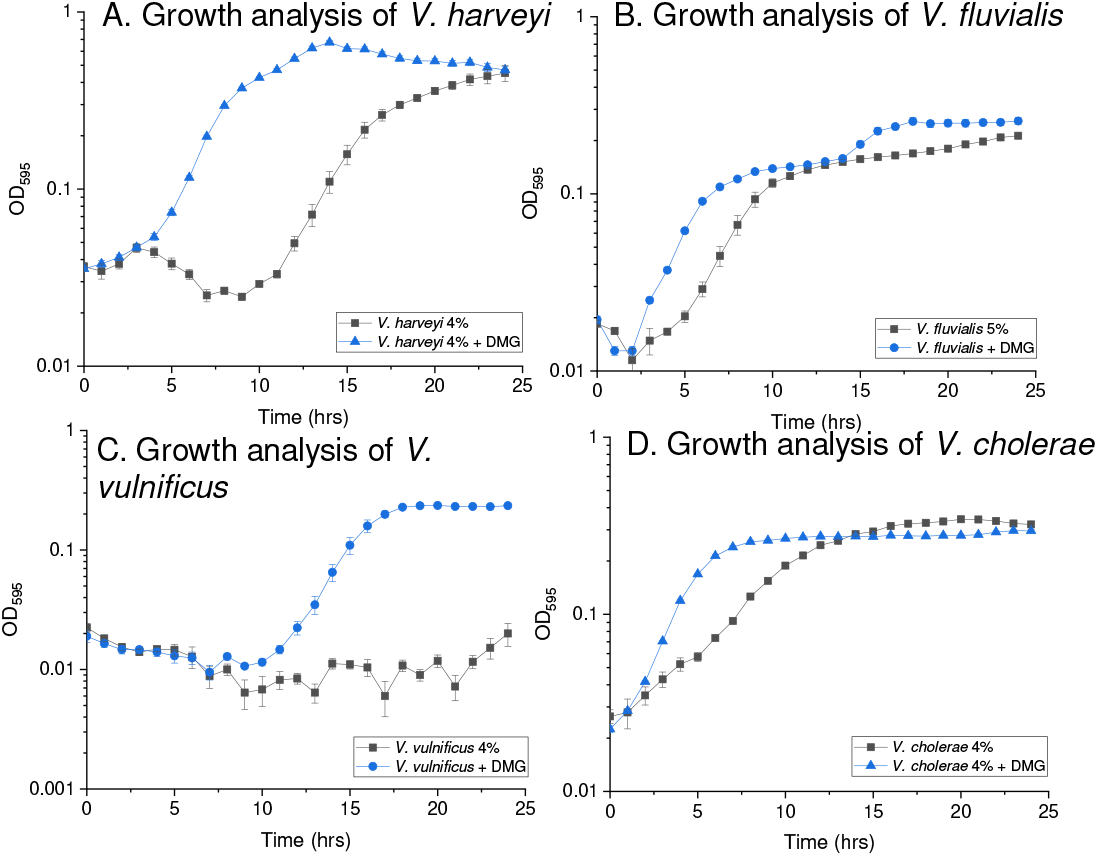
Growth analyses of **(A)** *V. harveyi* 393 and **(C)** *V. vulnificus* YJ016 **(D),** *V. cholerae* N16961, in M9G 4%NaCl with and without exogenous DMG. Growth analyses of **(B)** *V. fluvialis* growth analysis in M9G 5%NaCl with and without exogenous DMG. Optical density (OD_595_) was measured every hour for 24 hours. Mean and standard error of two biological replicates are shown.

Because DMG is an intermediate in glycine betaine catabolism, and many bacteria possess genes for the degradation of DMG to sarcosine, there is the possibility that DMG can be used as a carbon source in these species (40–42). Therefore, we examined whether any *Vibrio* species could utilize DMG as a sole carbon source by growing strains in M9 supplemented with DMG as the sole carbon source. None of the species tested could utilize DMG as a carbon source, demonstrating that DMG is a *bona fide* compatible solute for *Vibrio* species (**Fig. S2**).

### BCCTs are responsible for transport of DMG

DMG was identified as an alternative substrate of the *E. coli* transporter ProP, which is a member of the major facilitator superfamily (MFS) (43). In *Bacillus subtilis* DMG was transported into the cell as an omsolyte via an ABC-family transporter OpuA (44). We examined whether any of the BCCTs in *V. parahaemolyticus* were responsible for transport of DMG and to accomplish this we used a *bccT* null mutant (quadruple Δ*bccTI-bccT3-bccT4-bccT2*). We grew wild type and the *bccT* null mutant in M9G 6%NaCl supplemented with and without DMG at 37C for 24 h. In the wild-type strain, in the absence of DMG there was a 4 h lag phase whereas in the presence of DMG the lag phase was < 1 h and cells had a faster doubling time indicating that DMG is transported into the cell effectively (**Fig. 4**). However, the *bccT* null mutant did not exhibit a reduced lag phase or increased growth rate in the presence of DMG (**Fig. 4**), which indicated that a BCCT carrier is required for transport of DMG. Although, *V. parahaemolyticus* encodes two ABC-family compatible solute transporters, ProU1 and ProU2, which are presence in the *bccT* null mutant, these do not appear to be involved in DMG uptake.

**Figure 4.**
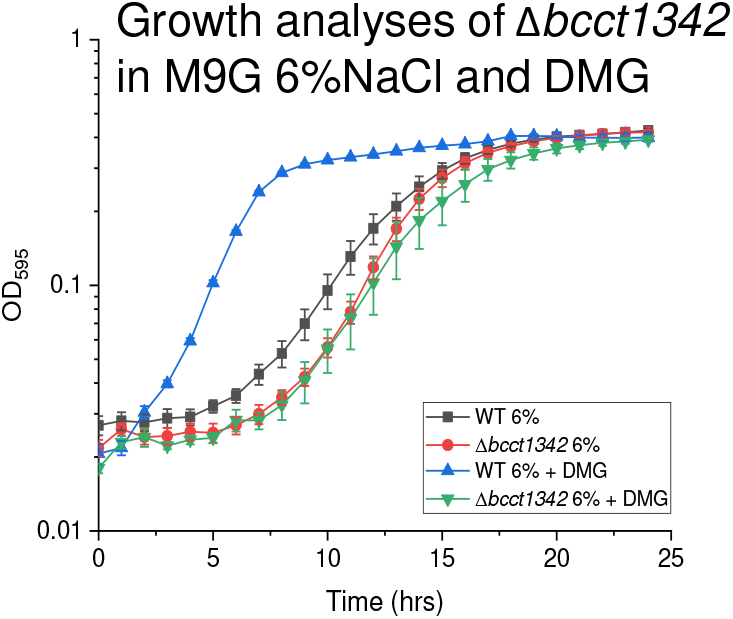
Growth analysis of wild type (WT) and *bccT* null (Δ*bccT1*-Δ*bccT3*-Δ*bccT4*-Δ*bccT2*) mutant in M9G 6%NaCl with and without DMG. Optical density (OD_595_) was measured every hour for 24 hours; mean and standard error of at least two biological replicates are displayed.

To determine which of the four BCCTs transports DMG, a set of four triple *bccT* mutants each possessing a single functional *bccT* was utilized in growth assays. The triple Δ*bccT2-bccT3-BccT4* mutant, which contains only *bccT1*, had a slightly reduced lag phase and a slightly faster growth rate through exponential phase in M6G 6%NaCl supplemented with DMG (**Fig. 5A**). This suggested that BccT1 transported DMG with low efficiency. In contrast, the triple Δ*bccT1-bccT3-bccT4* mutant, which contains only *bccT2*, had a similar reduction in lag phase and faster growth rate as wild type (**Fig. 5B**). This suggested that BccT2 transported DMG as efficiently as the wild-type strain. The triple Δ*bccT1-bccT2-bccT4* mutant, which contains only *bccT3*, grew similar to the bccT1 only strain, with a very slightly reduced lag phase and grew slightly better through exponential phase in the presence of DMG (**Fig. 5C**). This suggested that DMG is transported by BccT3 with low efficiency. The Δ*bccT123* mutant showed no difference in growth in the absence or presence of DMG, which indicated that BccT4 does not transport DMG into the cell (**Fig. 5D**).

**Figure 5.**
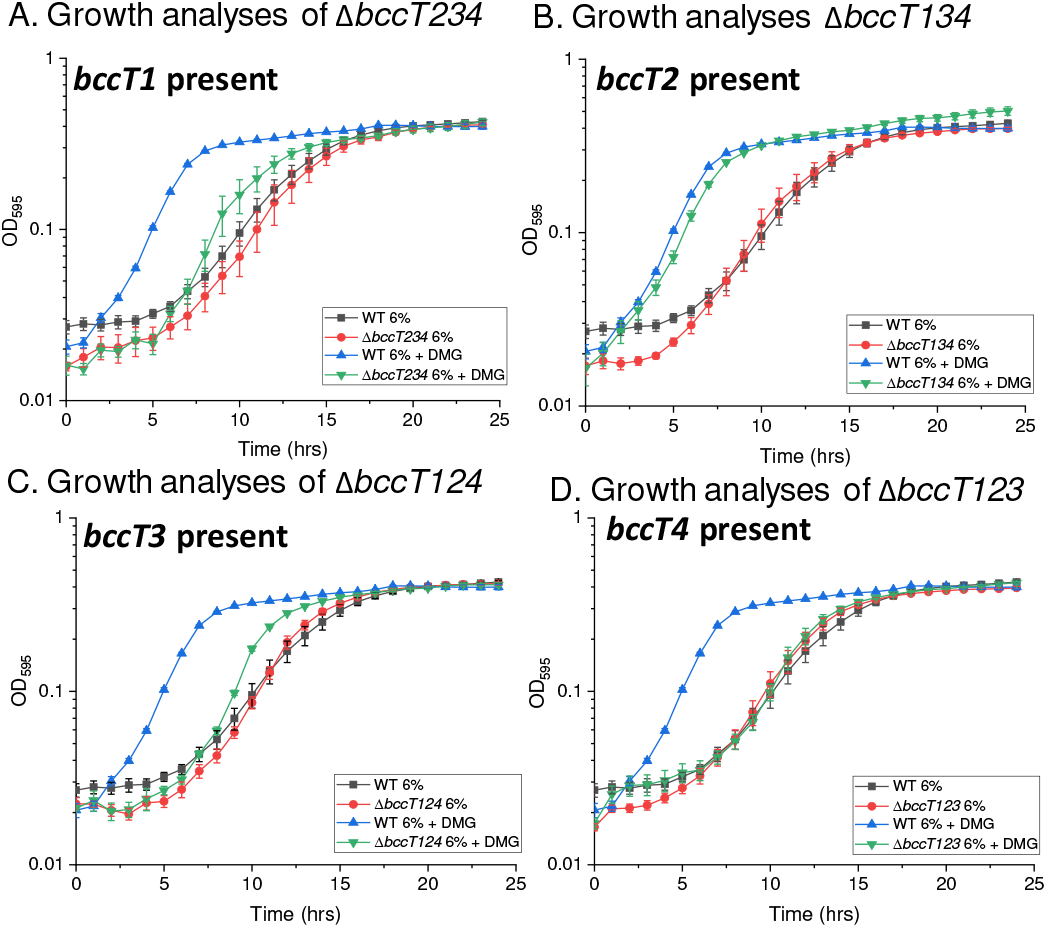
Growth analysis of wild type (WT) and triple mutants **(A)** Δ*bccT2*-Δ*bccT3*-Δ*bccT4*, **(B)** Δ*bccT1*-Δ*bccT3*-Δ*bccT4*, **(C)** Δ*bccT1*-Δ*bccT2*-Δ*bccT4*, or **(D)** Δ*bccT1*-Δ*bccT2*-Δ*bccT3* in M9G 6%NaCl with and without DMG. Optical density (OD_595_) was measured every hour for 24 hours; mean and standard error of at least two biological replicates are displayed.

In order to examine DMG uptake by the BCCT transporter further, we used *E. coli* MKH13, a mutant strain that does not contain osmolyte transporters or biosynthesis genes and cannot grow in M9G 4%NaCl. We cloned each of the *bccT* genes into an expression plasmid and used each construct to complement *E. coli* MKH13 and examined growth in M9G 4%NaCl supplemented with DMG (**Fig. 6**). *Escherichia coli* MKH13 strains complemented with *bccT1*, *bccT2* and *bccT3* grew in the presence of DMG whereas the *bccT4* complemented strain and the empty expression plasmid did not (**Fig. 6**). The *E. coli* MKH13 pBAVP1456 (*bccT1*) and pBAVP1723 (*bccT2*) strains grew significantly better than the strain with pBAVP1905 (*bccT3*) (**Fig. 6**). These indicated that BccT1 and BccT2 are more efficient transporters of DMG than BCCT3, and BCC4 is not a transporter of this substrate.

**Figure 6.**
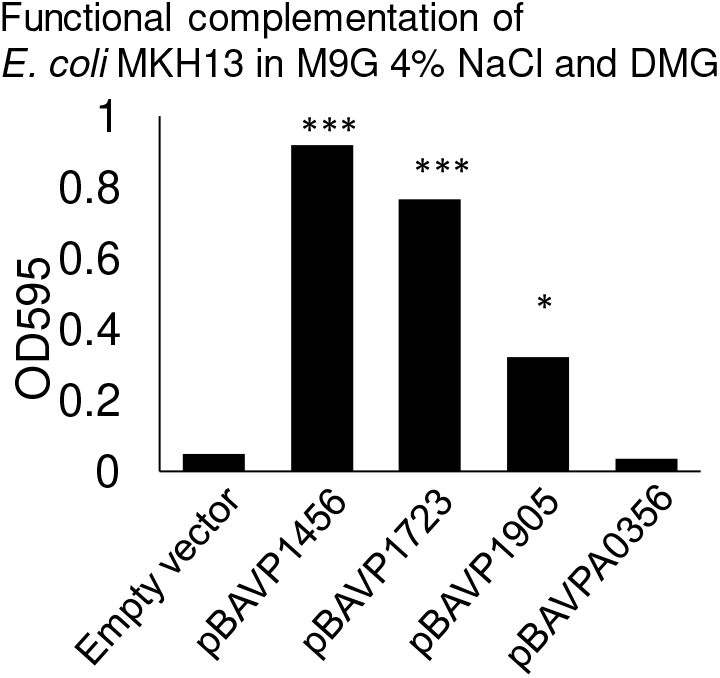
*E. coli* strain MKH13 was grown in M9G 4%NaCl and complemented with **(A)** *bccT1* (pBAVP1456), **(B)** *bccT2* (pBAVP1723), **(C)** *bccT3* (pBAVP1905), or **(D)** *bccT4* (pBAVPA0356), expressed from pBAD33. Strains were grown for 24 hours in the presence of DMG and the final optical density (OD_595_) was compared to that of a strain harboring empty pBAD33. Mean and standard error of at least two biological replicates are shown. Statistics were calculated using a Student’s t-test (*, P < 0.05;***, P < 0.001).

### BccT1 sequence homology to structurally characterized BCCTs

From a structure/function standpoint, BccT1 is of particular interest as it can transport glycine betaine, DMG, and ectoine, which we confirmed by demonstrating uptake in *E. coli* MKH13 pBAVP1456 (*bccT1*) (**Fig. 7**). While glycine betaine and DMG have similar structures with methylated head groups, ectoine is a cyclic compound and may require coordination by different amino acid residues in the transporter (22). Furthermore, transporters of ectoine typically do not possess the conserved aromatic residues located in TM4 and TM8 that coordinate trimethylammonium substrate binding such as glycine betaine (22). Therefore, first we compared BccT1 to structurally characterized BCCT proteins by hydropathy analysis. The hydropathy profile of BccT1 was aligned with that of BetP from *Corynebacterium glutamicum* (CgBetP), a glycine betaine transporter whose structure has been studied extensively. We found BccT1 possessed 12 TM segments (TM1 to TM12) along with N- and C-terminal tail extensions. BccT1 shared matched positions with 89% of residues in CgBetP, which indicated the structures were highly conserved (**Fig. S3**). In CgBetP, the conserved residues that form the glycine betaine binding pocket in TM4 are Trp189, Trp194, Tyr197 and one residue located in TM8, Trp374. An additional residue located in TM8 below the binding pocket (Trp377) is thought to be involved in substrate coordination during conformational changes (24). In addition, *C. glutamicum* BCCT-family transporters LcoP (CgLcoP) and EctP (CgEctP), have been reported to uptake glycine betaine and ectoine (45, 46). We aligned the protein sequences of each of these transporters with CgBetP and BccT1, in order to compare residues in each protein (**Fig. S4**). CgBetP, CgLcoP and BccT1 possessed identical amino acids to residues corresponding to TM4 Trp189, Trp194, Tyr197, and TM8 Trp374 and Trp377 suggesting that these proteins coordinate glycine betaine with the same residues (**Fig. S4**).

**Figure 7.**
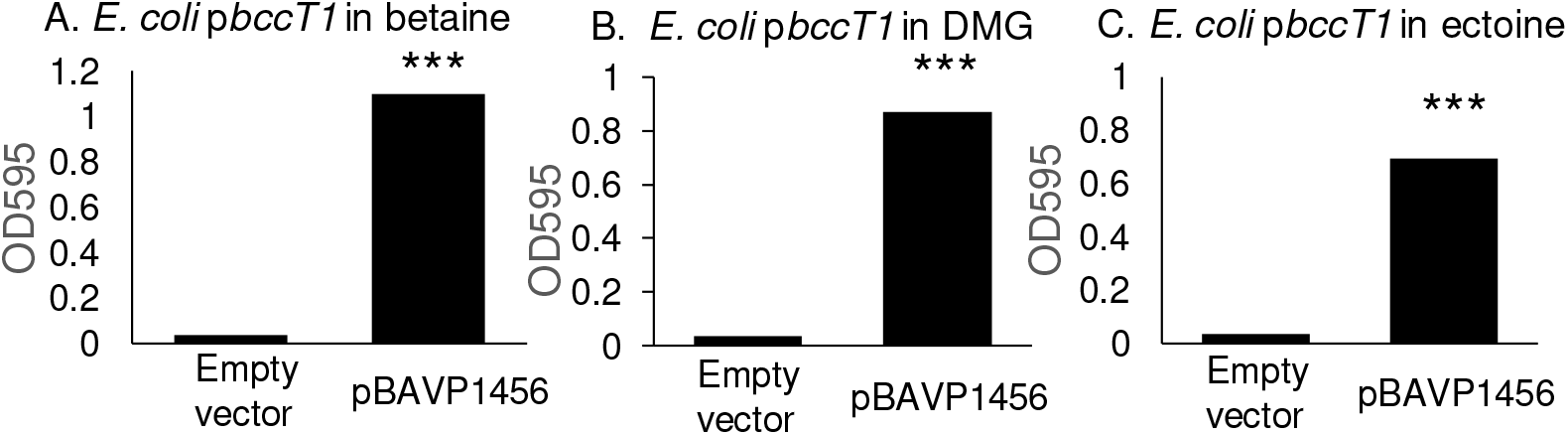
Complementation of *E. coli* MKH13 with *bccT1* (pBAVP1456) and growth in M9G 4%NaCl with **(A)** glycine betaine, **(B)** DMG, and **(C)** ectoine. Strains were grown for 24 hours and the final optical density (OD_595_) was compared to that of a strain harboring empty pBAD33. Mean and standard error of at least two biological replicates are shown. Statistics were calculated using a Student’s t-test (***, P < 0.001).

### Structural Modelling of BccT1

A BLASTP search of BccT1 sequence against Protein Data Bank (PDB) provided glycine betaine transporter CgBetP (PDB ID 4AIN) as the highest scoring hit with a sequence identity of 37% to BccT1 over 80% query coverage. CgBetP is an active trimer correctly represented in its three-dimensional structure with the three protomers (chains) showing different stages of substrate transport cycle with an alternating-access mechanism (47). More specifically, the chain A of CgBetP structure represents closed apo state (C_c_) whereas chains B and C represent a closed substrate-bound (C_c_S) and an open substrate-bound (C_i_S) state, respectively. The chain B and chain C of CgBetP were used as templates to generate C_c_S and C_i_S-like homology model of BccT1, respectively. The energy-minimized models of BccT1 were verified using a 3D profile method of Verify3D that evaluates the correlation between amino acid sequence (1D) and the model (3D) by comparing it to the other known structures in the database. The C_c_S and C_i_S model of BccT1 showed Verify3D scores of 88.6% and 84.0% respectively, which indicated good quality of the models (**Fig. S5**). The stereo-chemical properties of the models were examined by Ramachandran plot using PROCHECK. Both C_c_S and C_i_S BccT1 models contain 99.2% of all the residues in the allowed region and none in the disallowed region (**Fig. S6**).

Our structural models of BccT1 have cylindrical shapes and are composed of eleven trans-membrane helices (2–12) and a periplasmic helix (H7) (**Fig. 8A**). The first 108 amino acids of BccT1 could not be modeled because of their poor sequence similarity with CgBetP. BccT1 also lacks a cytosolic C-terminal domain (CTD), which is important for osmo-sensing in CgBetP (48). Nonetheless, BccT1 models are very similar to the CgBetP structure with an RMSDs of 0.26Å (399 Cα atoms) and 0.22 Å (391 Cα atoms) for C_c_S and C_i_S models, respectively. The residues involved in the central binding pocket, cytoplasmic and periplasmic gates as well as the rest of the substrate pathway are mostly conserved in BccT1. More specifically, a comparison of BccT1 models with the glycine betaine-bound structure of CgBetP showed that all the substrate binding residues are conserved and located in TM4 (Trp203, Trp208, Tyr211) and TM8 (Trp380, Trp381 and Trp384) (**Fig. S4**).

**Figure 8.**
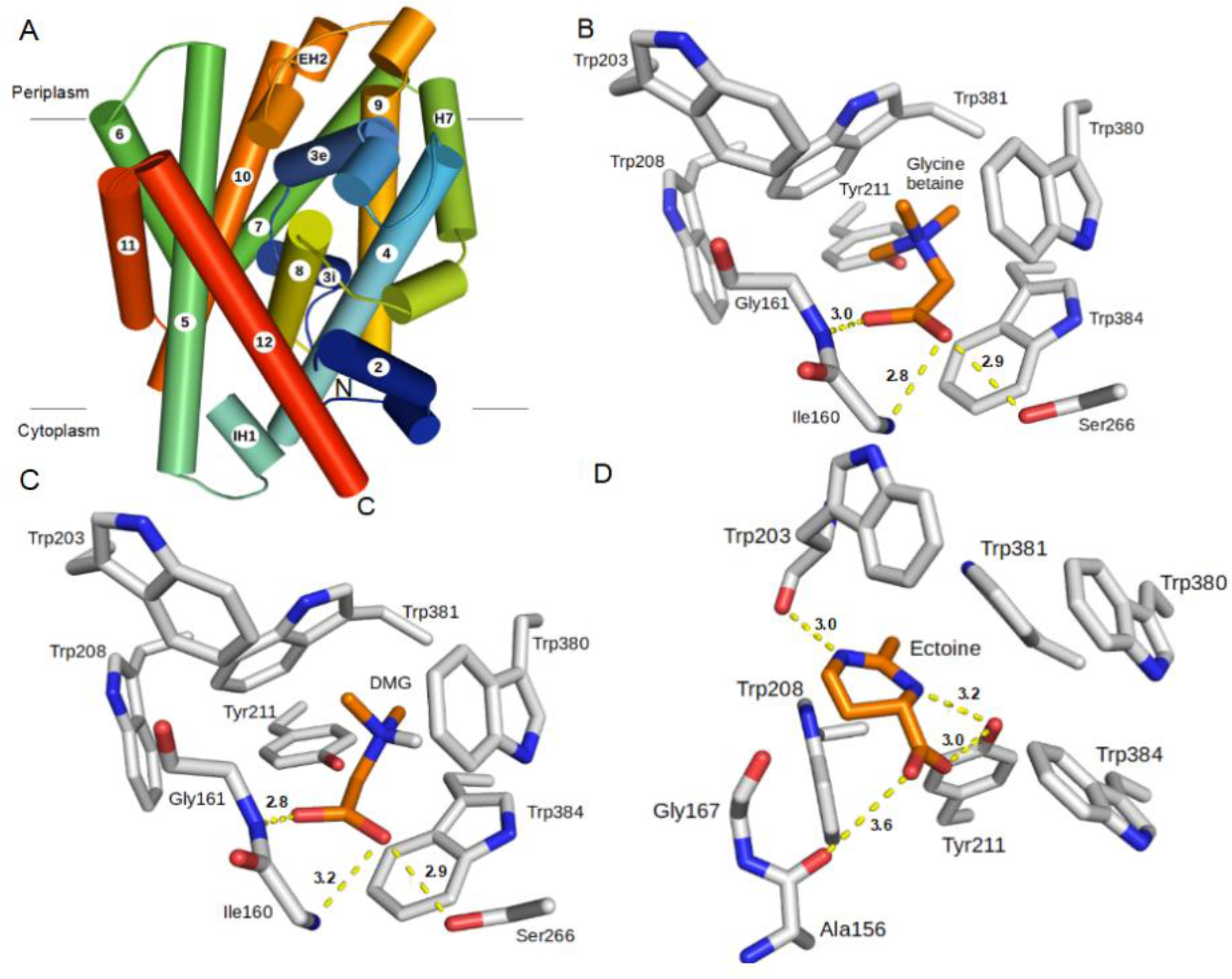
Predicted overall structure of BCCT1 and its active site interactions with different substrates. **(A)** A side view of the overall structure of BCCT1. α-helices are depicted by cylinders and directionality of the polypeptide chain is shown by a color gradient from blue (N-terminus) to red (C-terminus). Eleven trans-membrane helices (TM2-TM12) and a periplasmic helix (H7) make the core of the structure. Two small helices, IH1 and EH2, connect TM4-TM5 and TM9-TM10 respectively. **(B-D)** Interactions of the active site residues of BccT1 with the substrates. **(B)** Interacting residues of BccT1 C_c_S state with docked glycine betaine and **(C)** DMG; **(D)** interactions of BccT1 residues in a C_i_S state with docked ectoine. Bonds are depicted as sticks; substrates and BccT1 residues are colored orange and gray, respectively. Yellow dashed lines show hydrogen bonding distances between BccT1 residues and various atoms in the substrates. Illustrations are prepared using Pymol (The PyMOL Molecular Graphics System, Version 2.0 Schrödinger, LLC).

The closed state (C_c_S) represents a transition state between outward- and inward-facing open states where the substrate is bound in a central cavity with both substrate entry and exit ports occluded. Unlike open states, the substrate makes optimal interactions with the active site residues in the C_c_S state (47), therefore, C_c_S model of BccT1 was used for our substrate docking studies. The glycine betaine-docked BccT1 model showed binding free energy change (ΔG) of - 4.8 kcal/mole (**Table 1**). Briefly, the aromatic rings of Trp380, Trp381 and Trp384 form a small hydrophobic pocket that binds glycine betaine in this BccT1 model (**Table 1 and Fig. 8B**). The quaternary amine group of glycine betaine binds to this pocket by cation-π and van der Waals interactions. Other aromatic residues Trp203, Trp208 and Tyr211 of BccT1 also contribute to the hydrophobic interactions with glycine betaine. The glycine betaine carboxylic group interacts with the peptide backbones of Ile160 and Gly161 as well as side chain of Ser266 via hydrogen bonding (**Fig. 8B**).

**Table 1.** Results of the docking study showing free energy change upon substrate binding and a list of BccT1 residues mediating interactions with each substrate.

|  | **Glycine Betaine** | **DMG** | **Ectoine** |
| --- | --- | --- | --- |
| **Binding free energy (kcal/mol)** | -4.8 | -4.4 | -6.1 |
| **Hydrogen bonding interactions** | Ilu160, Gly161, Ser266 | Ilu160, Gly161, Ser266 | Trp203, Tyr211 |
| **Van der Waals interactions** | Trp203, Trp208, Tyr211, Trp380, Trp381, Trp384 | Trp203, Trp208, Tyr211, Trp380, Trp381, Trp384 | Trp208, Trp380, Trp381, Trp384 |

### Docking analysis of DMG and ectoine binding residues in BccT1

Since BccT1 is a unique BCCT-family transporter that can transport multiple osmolytes, we sought to determine the DMG and ectoine binding residues in BccT1 by docking. Docking of DMG to the BccT1 C_c_S model (**Fig. 8C**) predicted a mode of interaction identical to glycine betaine with a favorable binding free energy change (ΔG) of −4.4 kcal/mole (**Table 1**). This was expected since DMG and glycine betaine have very similar chemical structures.

To identify the mode of ectoine binding to BccT1, ectoine was first docked to the active site of the C_c_S model of BccT1. However, free energy change (ΔG) of ectoine binding to C_c_S model of BccT1 was too low (−2.8 kcal/mol), likely because of spatially constrained fitting of the large ectoine ring into the smaller hydrophobic pocket comprised of the aromatic rings of Trp380, Trp381 and Trp384. Therefore, we turned our attention to the C_i_S model of BccT1 ectoine docking, keeping all other docking parameters constant. The substrate binding site in the C_i_S state differs from C_c_S state in the relative orientations of the Trp380 and Trp381 side chains remodeling the hydrophobic pocket. More specifically, during the C_c_S to C_i_S transition, aromatic rings of Trp380 and Trp381 are known to flip 90 degrees, opening up the hydrophobic pocket (47). Interestingly, ectoine docking to the C_i_S state of BccT1 was accompanied by a significant free energy change (ΔG) of −6.1 kcal/mol (**Table 1**). The ectoine binding in this BccT1 state is stabilized by hydrogen bonding interactions between ectoine carboxyl group and the peptide backbone of Ala157 as well as the side chain of Tyr211 (**Fig. 8D**). Further, the ectoine ring nitrogens are involved in hydrogen bonding with Tyr211 and the peptide backbone of Trp203. The bound ectoine also shows van der Waals interactions with the aromatic residues Trp203, Trp208, Trp281 and Trp384 (**Fig. 8D**).

### Site directed mutagenesis of BccT1 uncovers ectoine binding pocket

The above-mentioned structural modeling and docking experiments predicted BccT1 residues that are likely involved in binding and transport of glycine betaine, DMG and ectoine. All these residues are conserved in CgBetP, and interestingly, mutations targeting four of the corresponding CgBetP residues, namely Trp189, Trp194, Tyr197, and Trp377, have been shown to significantly abrogate transporter function (24). Therefore, we selected the corresponding residues in BccT1 namely, Trp203, Trp208, Tyr211, and Trp384, for a mutagenesis study. We utilized functional complementation of *E. coli* MKH13 as a readout for uptake of glycine betaine, DMG and ectoine by BccT1 with single amino acid substitutions: Trp203Cys, Trp208Leu, Tyr211Leu and Trp384Leu. Amino acid substitutions corresponding to these positions in CgBetP, which conserve the bulky side chain of the amino acid being substituted, have been demonstrated to affect glycine betaine uptake in CgBetP (24). We also tested a BccT1 mutant with all four of these residues replaced by alanine, which is chemically inert and possesses a non-bulky methyl functional group. Replacement of Trp203 resulted in reduced growth of the *E. coli* MKH13 strain in the presence of glycine betaine, while strains harboring the other three single-replacement mutants Trp208Leu, Tyr211Leu, and Trp384Leu grew similarly to a strain harboring WT BccT1 (**Fig. 9A**). This indicates that Trp203 is important, but not essential, for uptake of glycine betaine by BccT1. In the absence of each of the other three residues 208, 211, or 384, glycine betaine may be accommodated by an alternate residue. Replacement of all four residues with alanine resulted in no growth of the *E. coli* MKH13 strain (**Fig. 9A**). This indicated that these four residues are involved in coordination of glycine betaine in BccT1.

**Figure 9.**
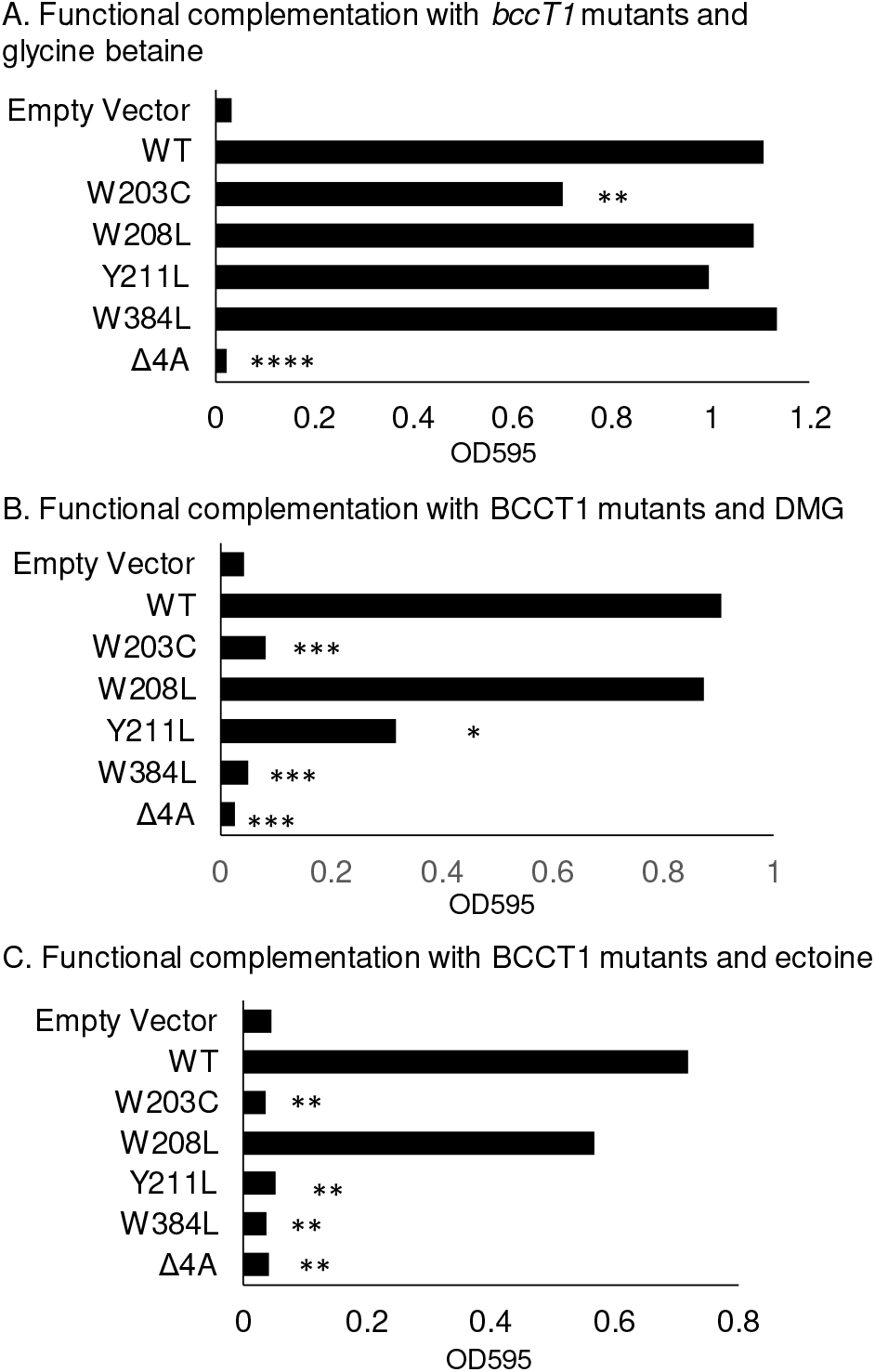
Functional complementation of *E. coli* MKH13 with *bccT1* (WT) and five *bccT1* mutants grown in **(A)** glycine betaine, **(B)** DMG, or **(C)** ectoine. Strains were grown for 24 hours and the final optical density (OD_595_) was compared to that that of the strain harboring wild type *bccT1* (pBAVP1456). Mean and standard error of at least two biological replicates are shown. Statistics were calculated using a Student’s t-test (*, P < 0.05; **, P < 0.01; ***, P < 0.001; ****, P < 0.0001).

Replacement of Trp203 or Trp384 resulted in no uptake of DMG, as strains harboring these mutants did not grow (**Fig. 9B**). Replacement of Tyr211 resulted in a reduced ability to uptake DMG, as evidenced by growth of this strain to a lower final OD (**Fig. 9B**). Replacement of all four residues resulted in no growth of *E. coli* MKH13, which indicated that these residues make up the binding pocket for DMG (**Fig. 9B**).

Replacement of residues Trp203, Tyr211 or Trp384 individually was sufficient to completely abolish uptake of ectoine by *E. coli* MKH13, as the strains harboring these mutants were unable to grow (**Fig. 9C**). However, replacement of Trp208 did not result in a statistically significantly difference in growth from a WT BccT1-expressing strain, indicating that Trp208 is not required for uptake of ectoine (**Fig. 9C**). Replacement of all four residues with alanine also resulted in abrogation of ectoine transport (**Fig. 9C**). Together these results indicated that DMG and ectoine share a binding pocket with glycine betaine. The coordination of DMG requires Trp203 and Trp384 while coordination of ectoine requires Trp203, Tyr211 and Trp384. These results suggest that coordination of DMG and ectoine cannot be as easily accommodated by alternate residues as can glycine betaine.

### Distribution of BccT1

The BccT1 protein is 562 amino acids and is present in all sequenced *V. parahaemolyticus* genomes (>800 genomes). A highly homologous protein is also present in all *V. alginolyticus, V. antiquarius, V. diabolicus* strains (95% sequence identity), *V. natriegens, V. nersis* (90% sequence identity), as well as *V. harveyi* and *V. campbellii* sequenced strains (86% sequence identity) (**Fig. S7**). In general, phylogenetic analysis indicated that BccT1 is conserved within the Harveyi clade and when present in other clades it is present in all strains of each species. Overall, within the family *Vibrionaceae*, BccT1 is present in 35 *Vibrio* species and in 11 *Photobacterium* species (**Fig. S7**). It is of interest to note that in all species, the residues corresponding to Trp 203, Trp 208, Tyr 211, and Trp 384, which coordinate substrates, are conserved (data not shown), suggesting an ability to uptake a range of osmolytes in these species.

## Discussion

Here we have demonstrated that *V. parahaemolyticus* can utilize additional compatible solutes DMG, GABA, TMAO, and creatine amongst others that have not been previously reported as osmolytes for Vibrio species. BccT1, BccT2 and BccT3 are capable of uptake of DMG, which is a highly effective osmoprotectant for *V. parahaemolyticus* and also is a potent osmoprotectant for *V. vulnificus*, *V. harveyi*, *V. cholerae* and *V. fluvialis*. DMG is the N-dimethyl derivate of glycine while glycine betaine is the N-trimethyl derivative. Some halophilic bacteria were shown to accumulate DMG, an intermediate compound produced during *de novo* biosynthesis (49–57). Additionally, DMG is an intermediate of aerobic glycine betaine catabolism, therefore should be available in the environment (40, 44, 58–60). DMG was found to be suitable for osmotic adaptation on its own, without modifications by bacteria (49). As stated previously, DMG is known to be transported by MFS family and ABC-family transporters (43, 44). Here, we demonstrated that the BCCT family can also uptake DMG.

BccT1 and BccT2 effectively transport DMG when expressed in a heterologous *E. coli* background (**Fig. 6**). However, in the native background, a *V. parahaemolyticus* strain expressing only BccT2 is fully rescued to wild-type levels by the presence of DMG while a strain expressing BccT1 is only partially rescued (**Fig. 5A and 5B**). For our growth analyses, cells are first grown in low salinity (1% NaCl) and then upshocked into high salinity (6% NaCl). We have shown previously that *bcct2* is not induced by high salinity and has a basal level of transcription in the cell, whereas *bcct1* expression is repressed in low salinity conditions (37, 38). Therefore, in a strain expressing only *bcct2*, expression is constitutively active in low salinity, which may allow for more rapid adaptation to high salinity conditions via uptake of DMG, resulting in a reduced lag phase. In *C. glutamicum*, the transporter EctP is not induced by external osmolarity and EctP transport capacity is maximal at low osmolarity and does not increase in high osmolarity conditions (46). EctP also showed a broad substrate spectrum with moderate to low affinity for these substrates, which, taken together, suggested that EctP acts as a rescue system that is available at low osmolarity, ensuring that cells can respond to osmotic stress (46). The same could be true of BccT2 in *V. parahaemolyticus*, which has a broad substrate range, including glycine betaine, choline, DMG, and proline and therefore could act as a rescue system to scavenge available compatible solutes in the event of osmotic stress.

DMG was not shown previously to be transported by the BCCT family of carriers and the amino acid residues important for uptake of DMG by a BCCT are unknown. Our *in silico* docking study showed that glycine betaine was coordinated by identical residues in the binding pocket of BccT1 (**Fig. 8B**) as have been previously reported for CgBetP (24, 25). Coordination of DMG by BccT1 was identical to that of glycine betaine (**Fig. 8C**). Likewise, an ATP-binding cassette (ABC) transporter, OpuAC, from *Bacillus subtilis* has been reported to interact very similarly with glycine betaine (PDB ID 2B4L) and a sulfonium analog of DMG, dimethylsulfonioacetate (DMSA) (PDB ID 3CHG) (61, 62). Our mutagenesis studies demonstrated that glycine betaine, DMG and ectoine are coordinated in the same binding pocket of BccT1, but the residues required for coordination are strict for DMG and ectoine, while glycine betaine may be accommodated by alternate residues in single amino acid mutants (**Fig. 9**). It was demonstrated previously in an ABC-type transporter, that while replacement of a single aromatic residue in the binding pocket results in decreased affinity of the protein for glycine betaine, the substrate is still coordinated with reasonable affinity. Only when any combination of two residues was mutated was transport completely abolished (63). We did not see a reduction in the ability of *E. coli* MKH13 to grow when only one residue is mutated, and uptake of glycine betaine was only completely abolished when all four residues were mutated. This is likely due to coordination of glycine betaine at alternate positions within the binding pocket that do not affect the overall growth ability of *E. coli* MKH13, but may affect the affinity of BccT1 for glycine betaine. Comparative protein analyses also demonstrated that in other *Vibrio* species with BccT1 homologs, the residues that coordinate glycine betaine, DMG and ectoine are conserved. There is a very high percent identity shared with BccT1 amongst BCCT-family transporters in these species and therefore the ability of these transporters to uptake a broad range of substrates is most likely conserved.

All of the substrates that *V. parahaemolyticus* was demonstrated to uptake are available in the marine environment, including glycine betaine, choline, DMG, and ectoine (4, 40, 58, 64). Five of the six compatible solute transporters encoded by *V. parahaemolyticus* are induced by high salinity, and collectively take up a broad range of compatible solutes (37, 38). Although there is redundancy in the compounds taken up by each BccT, the ability to uptake many different compatible solutes likely provides *V. parahaemolyticus* and other *Vibrio* species with a fitness advantage.

Members of the BCCT family are widespread among bacteria, present in both Gram-positive and Gram-negative bacteria as well as Archaea. For example, using the Interpro database (http://www.ebi.ac.uk/interpro/) and IPR000060 for BccT as a search, 23,000 BccT proteins fall within the domain Bacteria, 604 BccT proteins within the Archaea and 78 BccTs within the Eukaryota. Of the 604 Archaea representatives, 593 were from the Stenosarchaea group of which 549 were within *Halobacteria* suggesting an important function in osmotolerance. Surprisingly, there have been no studies on BCCT function from representatives of Archaea or Eukaryota to date.

## Methods

### Bacterial strains, media and culture conditions

All strains and plasmids used in this study are listed in Table 2. *Vibrio parahaemolyticus* strains were grown either in lysogeny broth (LB) (Fisher Scientific, Fair Lawn, NJ) with 3% (wt/vol) NaCl (LB3%) or M9 minimal media (47.8 mM Na_2_HPO_4_, 22mM KH_2_PO_4_, 18.7 mM NH4Cl, 8.6 mM NaCl; Sigma-Aldrich) supplemented with 2mM MgSO_4_, 0.1 mM CaCl_2_, 20 mM glucose as the sole carbon source (M9G) and NaCl (wt/vol), as indicated. Dimethylglycine (DMG) was used at a final concentration of 20 mM when supplied as a carbon source. *E. coli* strains were grown either in LB supplemented with 1% NaCl (LB1%) or M9G supplemented with 1% NaCl (M9G1%). All strains were grown at 37°C with aeration. Chloramphenicol (Cm), was added to the media at 25 μg/mL when necessary.

**Table 2.**
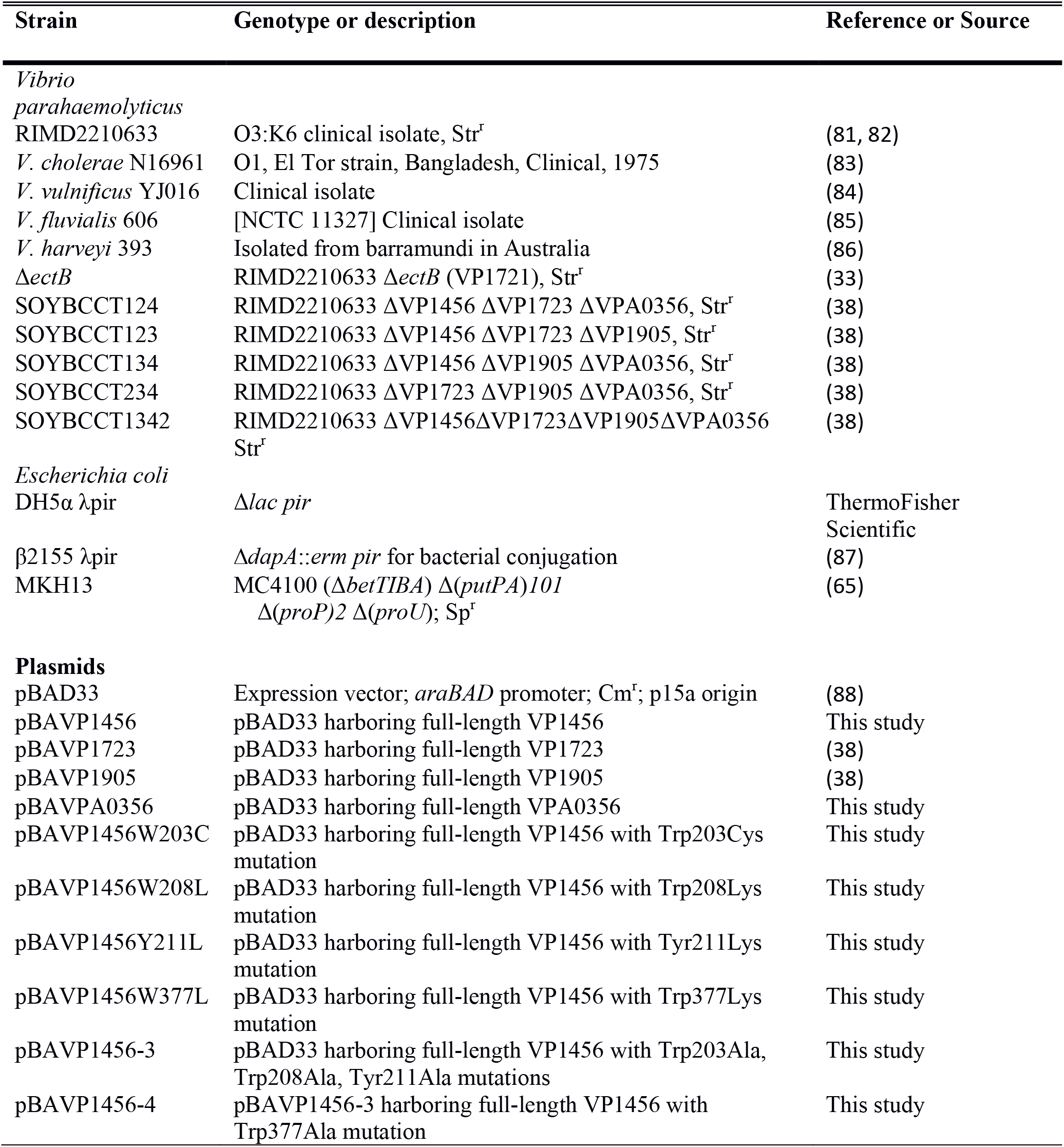
Strains and plasmids used in this study

### Growth analysis

*Vibrio parahaemolyticus* and an in-frame deletion mutant of *ectB* were grown overnight in M9 minimal media supplemented with 1% NaCl and 20 mM glucose as the sole carbon source. Cultures were subsequently diluted 1:50 into fresh medium and grown for five hours. Cultures were pelleted, washed two times with 1X PBS to remove excess salt, and then diluted 1:50 into M9G, and 100 uL was then added to each well of a 96-well Biolog PM9 plate containing different osmolytes and/or salt concentrations (Biolog, Inc., Hayward, CA). The plates were incubated at 37°C with intermittent shaking in a Tecan Sunrise microplate reader and OD_595_ was measured every hour for 24 hours. The area under the curve (AUC) was calculated using Origin 2018 for wild type (WT) and the Δ*ectB* mutant. Statistics were calculated using a Student’s t-test; growth in 6% NaCl in the presence of a compatible solute was compared to growth in 6% NaCl with no exogenous compatible solutes.

For growth analysis in individual compatible solutes, wild type (WT), Δ*ectB* mutant and quadruple Δ*bcct1*-Δ*bcct3*-Δ*bcct4*-Δ*bcct2* mutant (*bccT* null) strains were grown overnight in M9G 1%NaCl. Cultures were subsequently diluted 1:50 into fresh medium and grown for five hours to late exponential phase. Exponential cultures were then diluted 1:40 into 200 μL of M9G 6%NaCl medium with and without exogenous compatible solutes in a 96-well microplate and grown at 37°C with intermittent shaking for 24 hours. Compatible solutes N-N dimethylglycine (DMG), trimethylamine-N-oxide (TMAO) and γ-amino-N-butyric acid (GABA) were added to a final concentration of 500 μM. Growth analysis in each compatible solute was repeated following the above procedure with each of four triple *bccT* mutants, Δ*bccT2*-Δ*bccT3*-Δ*bccT4*, Δ*bcct1*-Δ*bccT3*-Δ*bccT4*, Δ*bcct1*-Δ*bccT2*-Δ*bccT4*, Δ*bcct1*-Δ*bccT2*-Δ*bccT3*.

Growth analyses of *V. cholerae*, *V. harveyi*, *V. vulnificus*, and *V. fluvialis* were conducted by growing strains overnight in M9G supplemented with 2% NaCl (M9G 2% NaCl). Strains were diluted 1:40 into 200 μL of M9G 4%NaCl medium, or M9G 5%NaCl for *V. fluvialis*, with and without exogenous DMG, in a 96-well microplate and grown at 37°C with intermittent shaking for 24 hours. To test DMG as a carbon source, *V. cholerae*, *V. vulnificus*, *V. fluvialis* and *V. parahaemolyticus* were grown overnight in LB1%; *V. harveyi* was grown in LB2%. Cells were pelleted, washed two times with 1 X PBS, and diluted 1:40 into 200 μL of M9 with 20 mM DMG as the sole carbon source and 1% NaCl (2% NaCl for *V. harveyi*). Strains were grown in M9G1% NaCl (2% NaCl for *V. harveyi*) as a control. Strains were grown in a 96-well microplate as described above.

### Functional complementation of *E. coli* strain MKH13 with VP1456, VP1723, VP1905, and VPA0356

Full-length VP1456 (*bccT1*) or VPA0356 (*bccT4*) were amplified from the *V. parahaemolyticus* RIMD2210633 genome using primers listed in Table 3. Gibson assembly protocol using NEBuilder HiFi DNA Assembly Master Mix (New England Biolabs, Ipswich, MA) was followed to ligate the VP1456 or the VPA0356 fragment with the expression vector pBAD33, which had been linearized with SacI. Regions of complementarity for Gibson assembly are indicated by lowercase letters in the primer sequence in Table 3. The resulting expression plasmids, pBAVP1456 or pBAVPA0356, were transformed into *E. coli* Dh5α for propagation. Plasmids were then purified, sequenced, and subsequently transformed into *E. coli* strain MKH13, which has large deletions that include all compatible solute transporters (*putP*, *proP*, *proU*) and the choline uptake and glycine betaine biosynthesis loci (*betT*-*betIBA*) (65). *E. coli* MKH13 strains containing pBAVP1456, pBAVP1723, pBAVP1905 or pBAVPA0356 were grown overnight in minimal media supplemented with 1% NaCl and 20 mM glucose (M9G1%) and subsequently diluted 1:40 into M9G supplemented with 4% NaCl (M9G 4%NaCl). *E. coli* MKH13 strains containing pBAVP1456, pBAVP1723, pBAVP1905 or pBAVPA0356 were grown overnight in M9G 1%NaCl with chloramphenicol and subsequently diluted 1:100 into M9G 4% NaCl and 500 μM of the indicated compatible solute and chloramphenicol for plasmid maintenance. Expression of each BccT was induced with 0.01% arabinose and functional complementation was determined by measuring OD_595_ after 24 hours growth at 37°C with aeration. Growth was compared to that of an MKH13 strain harboring empty pBAD33, which cannot grow in M9G 4%NaCl without exogenous compatible solutes. Statistics were calculated using a Student’s t-test.

**Table 3.**
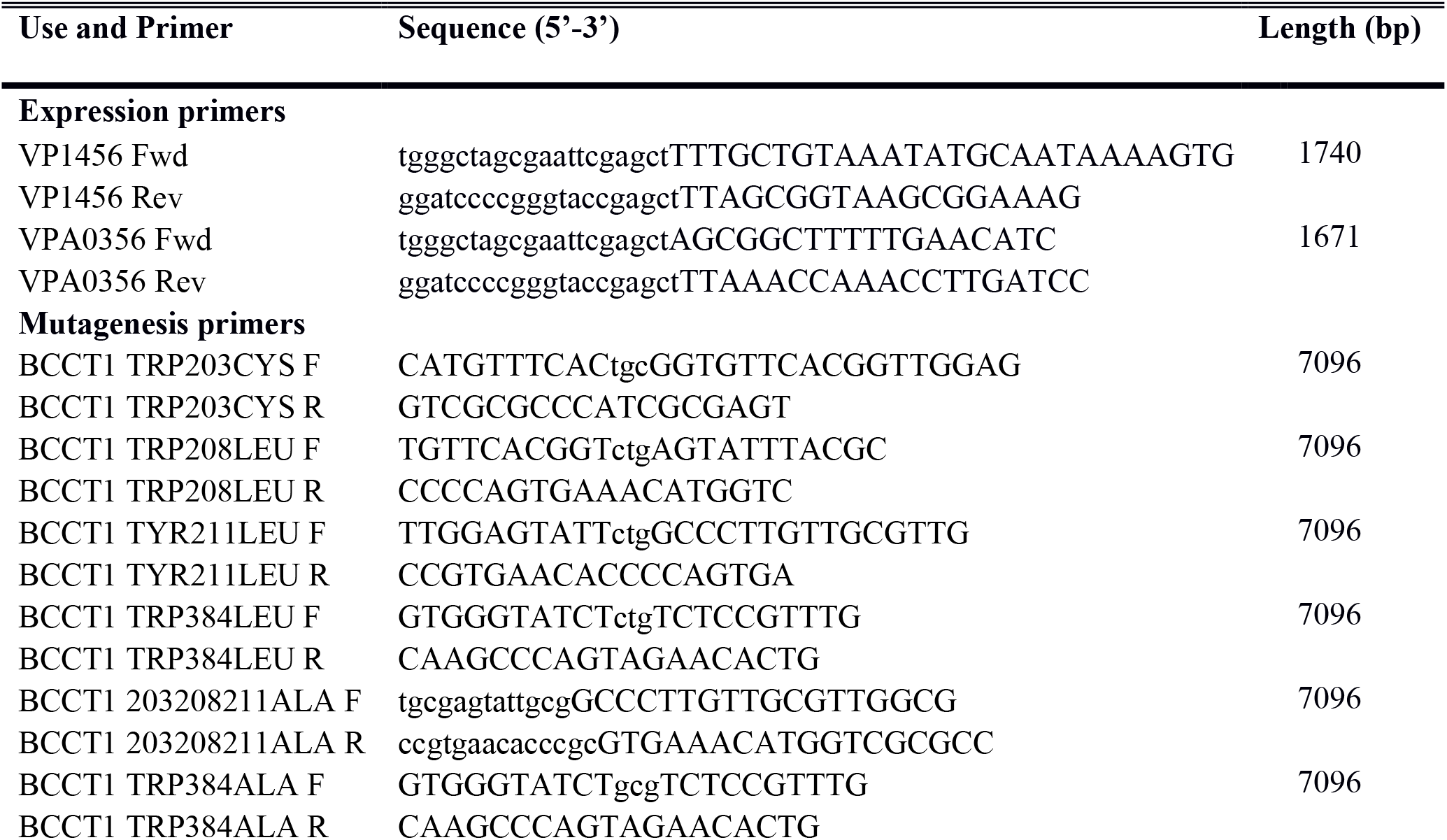
Primers used in this study

Site-directed mutagenesis was performed on pBPVP1456 with a Q5 site-directed mutagenesis kit (New England Biolabs, Ipswich, MA) and primers listed in Table 3. Primers were designed to create the nucleotide substitutions resulting in the following amino acid changes: Trp203Cys, Trp203Ala, Trp208Leu, Trp208Ala, Tyr211Leu, Tyr211Ala, Trp384Leu, and Trp384Ala. Site-directed mutagenesis was performed following the manufacturers’ protocol to create single amino acid substitutions Trp203Cys, Trp208Leu, Tyr211Leu, Trp384Leu.

Residues 203, 208, and 211 were mutagenized to encode for alanines, and residue 384 was subsequently mutagenized in this plasmid backbone to encode for an alanine.

### Homology modeling of BCCT1 and docking of glycine betaine, DMG and ectoine

BLAST search of BccT1 sequence against Protein Data Bank (PDB) as the search database showed highest sequence identity with the 3.2 Å X-ray crystal structure of Glycine Betaine transporter from *C. glutamicum* (CgBetP, PDB ID: 4AIN). Homology modeling of BccT1 was done in SWISS-MODEL server using CgBetP structure (PDB ID: 4AIN) as a template (66). CgBetP exists as a trimer in the structure with each protomer showing different states of substrate transport. Two different models resembling protomers from chain B and chain C of CgBetP structure, and therefore representing two different substrate binding states, were treated separately after modeling. Residues showing poor stereo-chemical properties or close-contacts in each monomeric model were fixed manually in COOT (67). Models were then subjected to energy minimization using 3Drefine server to reduce any other structural restraints (68). The quality of the resulting models was finally verified with Verify3D and PROCHECK (69, 70).

For docking studies, ligand models for glycine betaine, DMG and ectoine were obtained from Chemical Entities of Biological Interest (ChEBI) EMBL (71). Polar hydrogens were added to the ligand structures in PRODRG (72). The AutoDoc tools v1.5.6 were used to assign the rotatable bonds in the ligands and to add all polar hydrogens in the BccT1 models to prepare them for the docking. The ligand-centered maps for BccT1 models were assigned a grid size of 20 x 20 x 20 Å^3^. The docking experiments of glycine betaine, DMG and ectoine ligands to the BccT1 receptor model were performed using AutoDock Vina (73). The binding free energies and BccT1 residues making interactions with the ligands are listed in Table 1. The structural illustrations were prepared using PyMOL (The PyMOL Molecular Graphics System, Version 2.0 Schrödinger, LLC).

### Bioinformatics and phylogenetic analyses

Transmembrane helix probabilities of *C. glutamicum* BetP (CAA63771.1) and BccT1 (Q87PP5.1) were generated using OCTOPUS and aligned via the AlignMe program (http://www.bioinfo.mpg.de/AlignMe) (74–76). The *V. parahaemolyticus* protein BccTI (Q87PP5.1), and CgBetP (CAA63771.1), EctP (CAA04760.1), and LcoP (ASW14702.1) were downloaded from NCBI and aligned using the ClustalW algorithm (77). Aligned sequences were displayed and annotated using ESPript (http://espript.ibcp.fr/ESPript/cgi-bin/ESPript.cgi) (78).

Phylogenetic analysis was conducted using BccT1 (VP1456) protein as a seed to identity all homologs within the family *Vibrionaceae* with completed genome sequences available. Unique protein sequences that had >95% sequence coverage and >70% amino acid identity with BccT1 were obtained from NCBI database and aligned using the ClustalW algorithm (77). The evolutionary history of BccT1 was inferred by using the Maximum Likelihood method and Le_Gascuel_2008 model as determine by best fit model selection in MEGAX (79, 80). The tree with the highest log likelihood (−10453.23) is shown. The percentage of trees in which the associated taxa clustered together is shown next to the branches. Initial tree(s) for the heuristic search were obtained automatically by applying Neighbor-Join and BioNJ algorithms to a matrix of pairwise distances estimated using a Jones-Taylor-Thornton (JTT) model, and then selecting the topology with superior log likelihood value. A discrete Gamma distribution was used to model evolutionary rate differences among sites (5 categories (+*G*, parameter = 0.4143)). The tree is drawn to scale, with branch lengths measured in the number of substitutions per site. This analysis involved 49 amino acid sequences and a total of 525 positions in the final dataset.

## Supporting information

Supplemntary Figues S1-S7

## ACKNOWLEDGEMENTS

This research was supported by a National Science Foundation grant (award IOS-1656688) to E.F.B. and National Institutes of Health grant (award R35GM119504) to V.P. G.J.G. was funded in part by a University of Delaware graduate fellowship award. We thank members of the Boyd Group for constructive feedback on the manuscript.

---

**Figure S1.** Area under the curve analysis of growth of an Δ*ectB* deletion mutant with a subset of 23 osmolytes from the Biolog phenotypic microarray PM9 plate. Growth in the presence of individual osmolytes is compared to growth in minimal medium with 6% NaCl only. Mean and standard error of at least two biological replicates are shown. Statistics were calculated using a Student’s t-test (*, P < 0.05; **, P < 0.01).

**Figure S2.** Growth analyses of *V. parahaemolyticus* RIMD2210633 (down triangles), *V. harveyi* 393 (up triangles), *V. vulnificus* YJ016 (diamonds), *V. cholerae* N16961 (squares), and *V*. *fluvialis* (circles), in M9 with DMG (open shapes) as the sole carbon source or M9G (solid shapes). Optical density (OD_595_) was measured every hour for 24 hours. Mean and standard error of two biological replicates are shown.

**Figure S3.** The transmembrane helix probability of *V. parahaemolyticus* BccT1 was generated and aligned with *Corynebacterium glutamicum* BetP using AlignMe (http://www.bioinfo.mpg.de/AlignMe). Values close to 1 indicate a high probability of that sequence being in the membrane while 0 is a low probability of that sequence being in the membrane. Dots below the plot indicate gaps introduced during alignment.

**Figure S4.** The *V. parahaemolyticus* BCCT protein BccT1 was aligned with *Corynebacterium glutamicum* transporters BetP, LcoP and EctP and displayed using ESPript. Residues highlighted in red are strictly conserved. Residues highlighted in magenta are conserved in sodium-symporters. Residues marked with a cyan triangle have been demonstrated to be important for glycine betaine binding; residues highlighted in cyan are conserved. A green star denotes residues thought to be important for additional substrate binding; conserved residues are highlighted in green.

**Figure S5.** Verification of the overall quality of our predicted BccT1 models with Verify3D. (A) 88.6% of BccT1 residues in the C_c_S model have a 3D-1D score ≥0.2, (B) 84.0% of BccT1 residues in the C_i_S model have 3D-1D score ≥0.2.

**Figure S6.** Ramachandran plots verifying stereochemical properties of our BccT1 models. (A) C_c_S model of BccT1 has 90.2% residues in most favored, 9.0% in additionally allowed, and no residues in outlier regions (B) C_i_S model of BccT1 contains 91.5% residues in most favored, 7.7% in additionally allowed, and no residues in outlier regions.

**Figure S7.** Phylogenetic tree was inferred by using the Maximum Likelihood method and Le_Gascuel_2008 model in MEGAX. The tree with the highest log likelihood (−10453.23) is shown. The percentage of trees in which the associated taxa clustered together is shown next to the branches. This analysis involved 49 amino acid sequences and a total of 525 positions.

