## Supplementary material for "Investigations of dimethylglycine (DMG), glycine betaine and ectoine uptake by a BCCT family transporter with broad substrate specificity in *Vibrio* species": Supplemntary Figues S1-S7

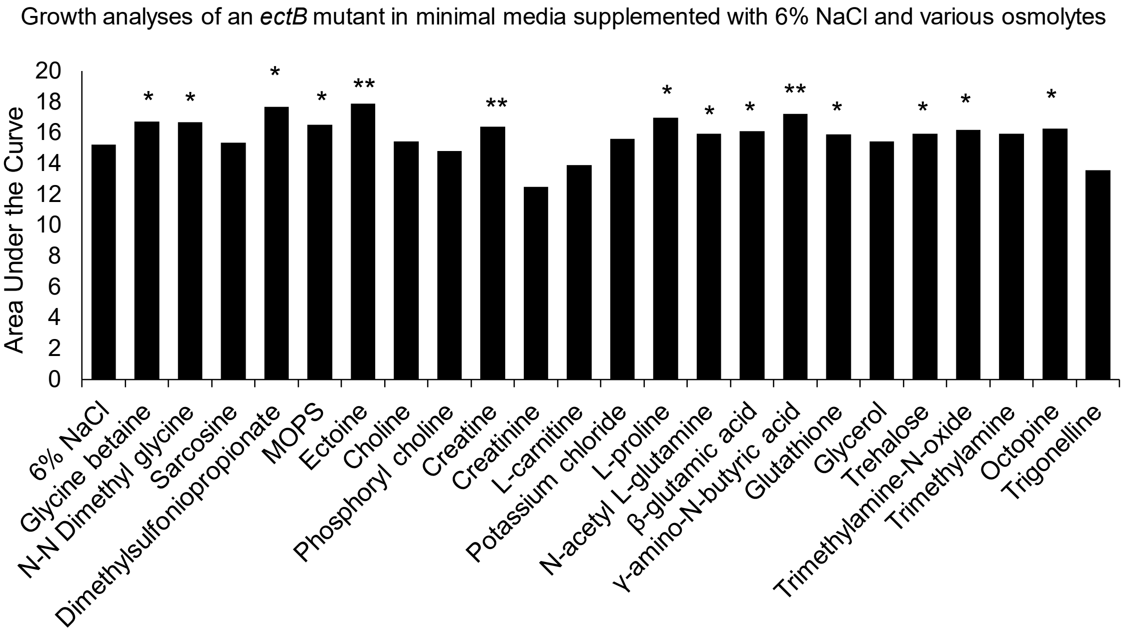


**Figure S1.** Area under the curve analysis of growth of an *ectB* deletion mutant with a subset of 23 osmolytes from the Biolog phenotypic microarray PM9 plate. Growth in the presence of individual osmolytes is compared to growth in minimal medium with 6% NaCl only. Mean and standard error of at least two biological replicates are shown. Statistics were calculated using a Student's t-test ((*, $P < 0.05$; **, $P < 0.01$).

Fig. S1

Growth analysis of *Vibrio* in M9G 1% NaCl or M9 DMG 1%NaCl

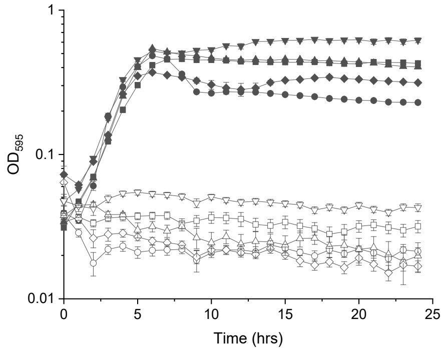


**Figure S2.** Growth analyses of *V. parahaemolyticus* RIMD2210633 (down triangles), *V. harveyi* 393 (up triangles), *V. vulnificus* YJ016 (diamonds), *V. cholerae* N16961 (squares), and *V. fluvialis* (circles), in M9 with DMG (open shapes) as the sole carbon source or M9G (solid shapes). Optical density ($OD_{595}$) was measured every hour for 24 hours. Mean and standard error of two biological replicates are shown.

Fig. S2

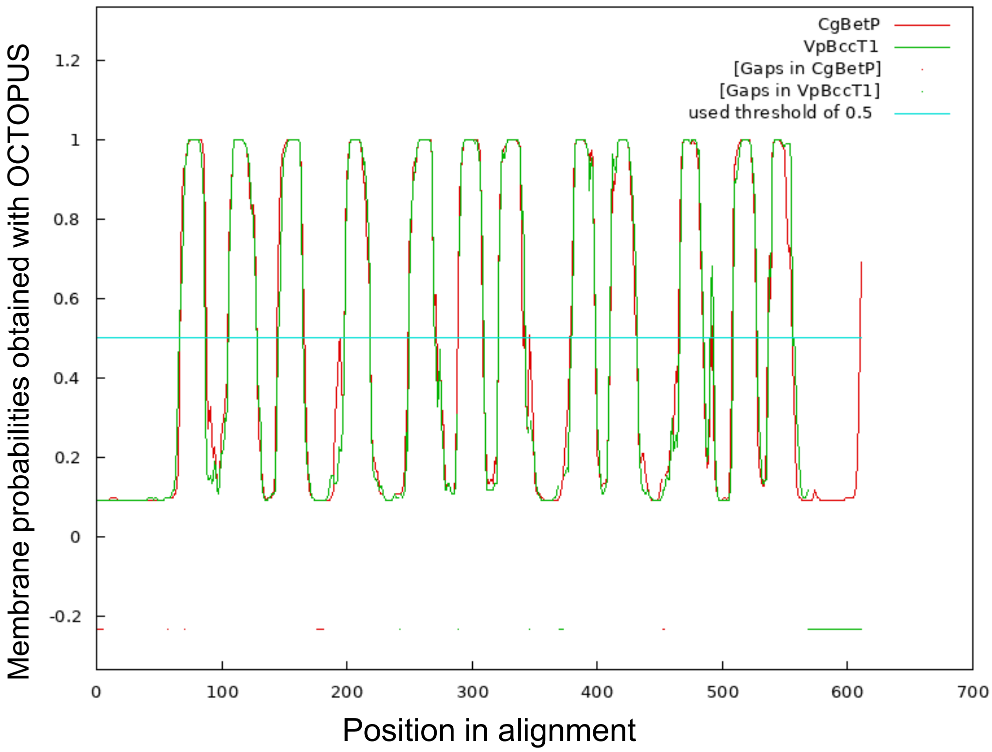


**Figure S3.** The transmembrane helix probability of *V. parahaemolyticus* BccT1 was generated and aligned with *Corynebacterium glutamicum* BetP using AlignMe (http://www.bioinfo.mpg.de/AlignMe). Values close to 1 indicate a high probability of that sequence being in the membrane while 0 is a low probability of that sequence being in the membrane. Dots below the plot indicate gaps introduced during alignment.

Fig. S3

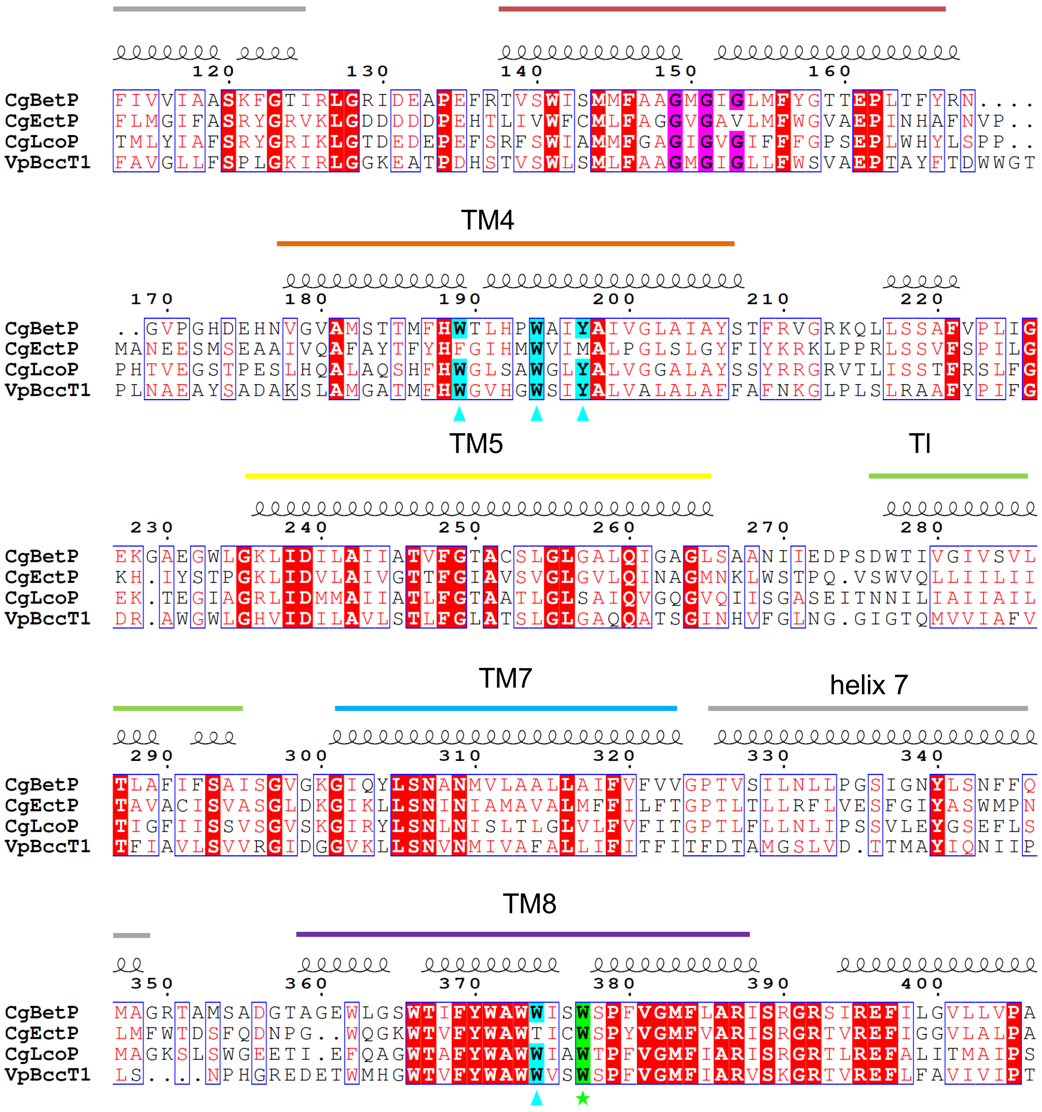


**Figure S4.** The *V. parahaemolyticus* BCCT protein VP1456 was aligned with *Corynebacterium glutamicum* transporters BetP, LcoP and EctP and displayed using ESPript. Residues highlighted in red are strictly conserved. Residues highlighted in magenta are conserved in sodium-symporters. Residues marked with a cyan triangle have been demonstrated to be important for glycine betaine binding; residues highlighted in cyan are conserved. A green star denotes residues thought to be important for additional substrate binding; conserved residues are highlighted in green.

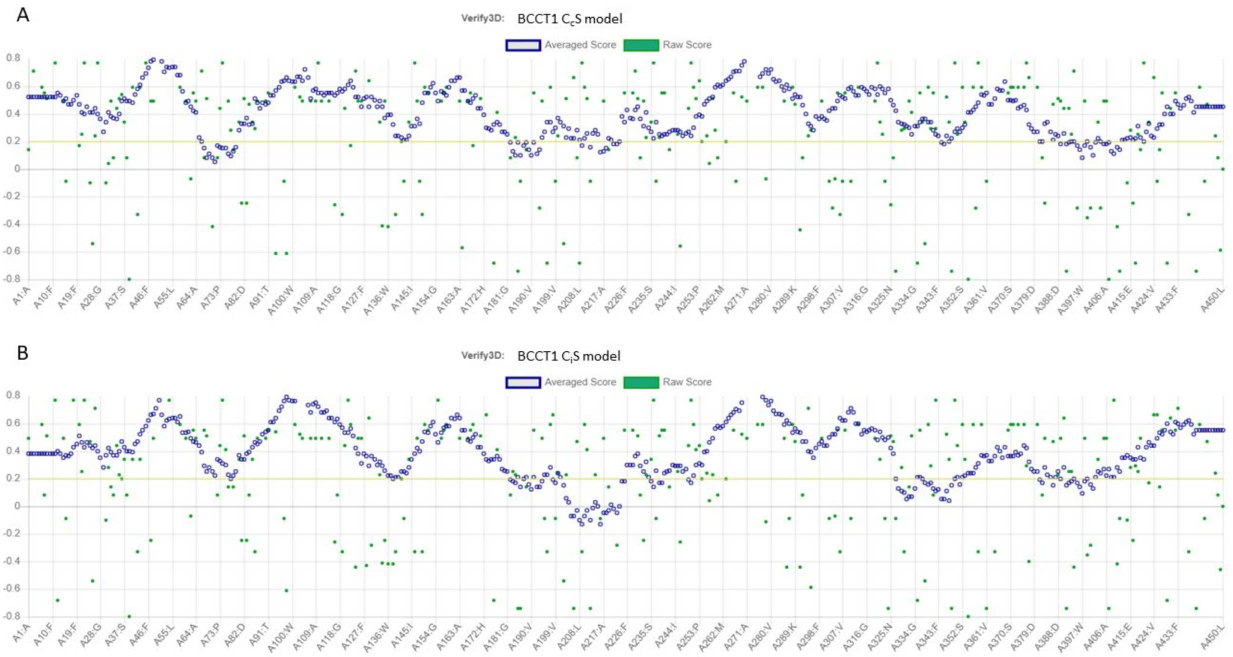


**Figure S5.** Verification of the overall quality of our predicted BccT1 models with Verify3D. (A) 88.6% of BccT1 residues in the $C_c$S model have a 3D-1D score ≥0.2, (B) 84.0% of BccT1 residues in the $C_i$S model have 3D-1D score ≥0.2.

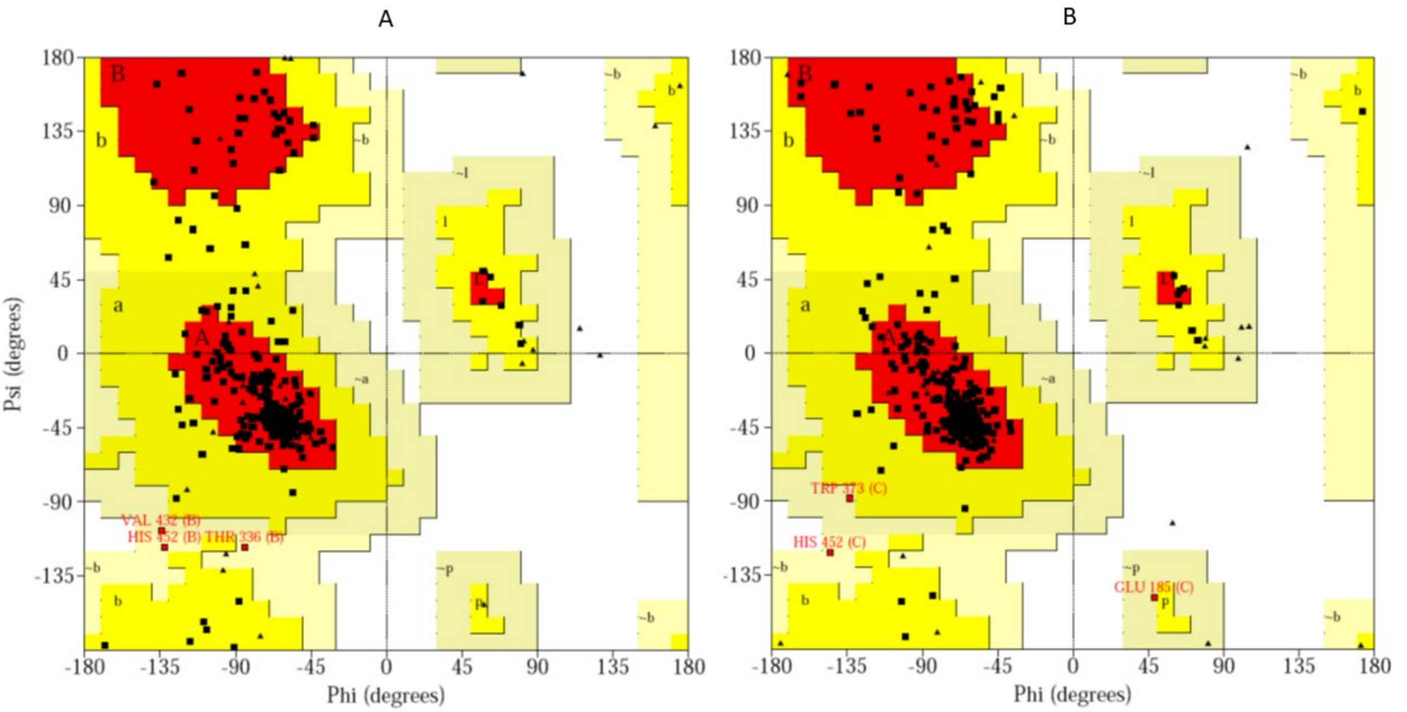


**Figure S6.** Ramachandran plots verifying stereochemical properties of our BccT1 models. (A) $C_cS$ model of BccT1 has 90.2% residues in most favored, 9.0% in additionally allowed, and no residues in outlier regions (B) $C_iS$ model of BccT1 contains 91.5% residues in most favored, 7.7% in additionally allowed, and no residues in outlier regions.

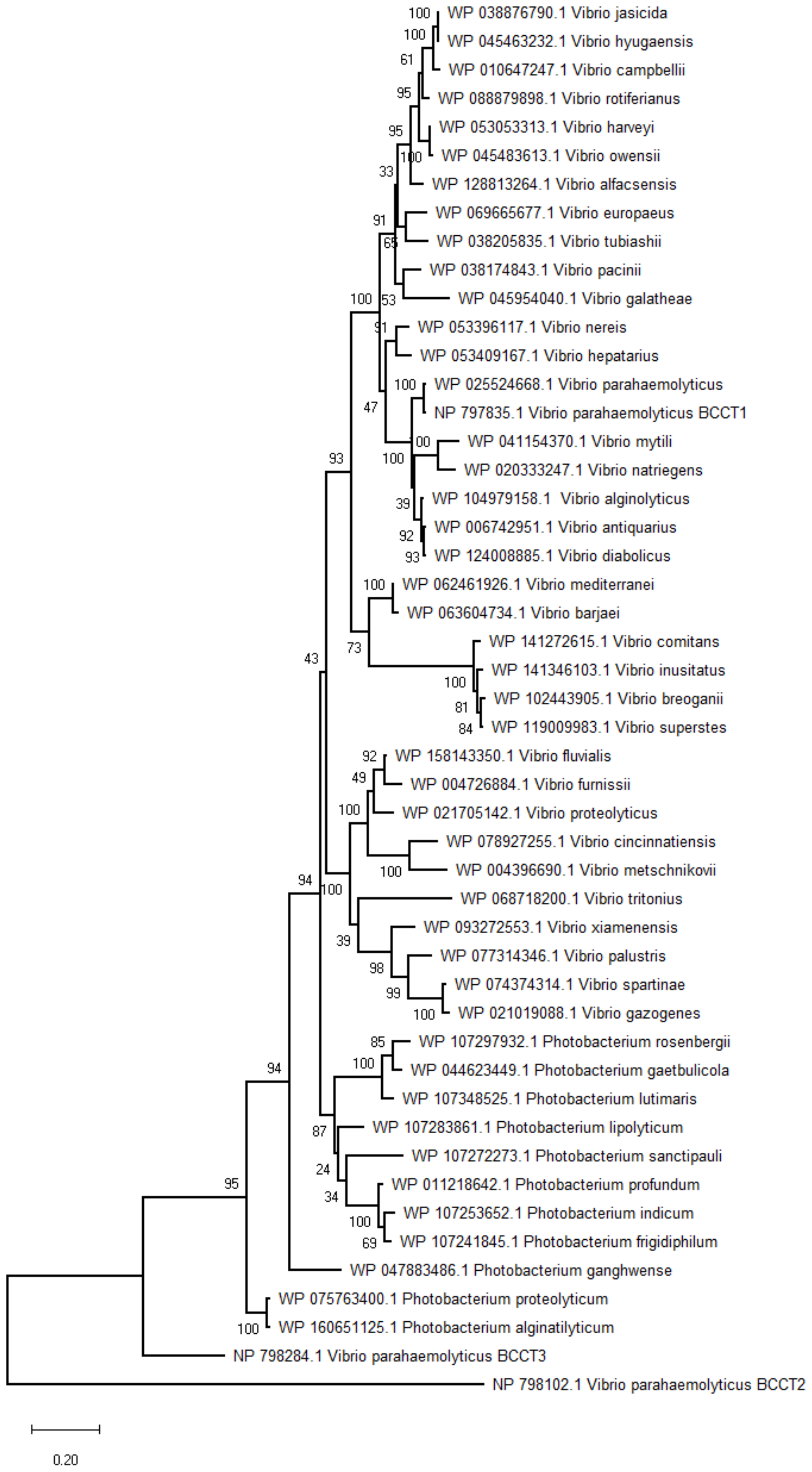


**Figure S7.** Phylogenetic tree was inferred by using the Maximum Likelihood method and Le_Gascuel_2008 model in MEGAX. The tree with the highest log likelihood (-10453.23) is shown. The percentage of trees in which the associated taxa clustered together is shown next to the branches. This analysis involved 49 amino acid sequences and a total of 525 positions.
